# Synchrotron diffraction reveals distinct collagen-associated distributions of EGFR-targeted antibody and affibody in pancreatic tumors

**DOI:** 10.64898/2026.11.10.765788

**Authors:** Cody C. Rounds, Sanzida Haque, Chengyue Li, Devin Nissen, Dhruv Ranjan, Marcella Vaicik, Rama S. Madhurapantula, Thomas C. Irving, Kenneth M. Tichauer

**Author notes:** Corresponding author: Kenneth M. Tichauer.

## Abstract

Tumor extracellular matrix may influence the distribution of targeted agents differently across molecular formats. We compared how an antibody and an affibody targeting the same receptor distribute relative to collagen-associated tissue structure in pancreatic tumors.

**Methods:** Cetuximab-IRDye700DX, an epidermal growth factor receptor (EGFR)-targeted fluorescent antibody, and ABY-029, an EGFR-targeted fluorescent affibody, were co-administered intravenously to 12 athymic nude mice bearing orthotopic pancreatic tumors: eight AsPC-1 and four PANC-1. Tumors were harvested at 3 or 24 h after injection, with four AsPC-1 and two PANC-1 mice at each time. Fluorescence images from 21 tissue slices were registered to scanning synchrotron X-ray diffraction (XRD) maps to compare each agent’s distribution with local collagen-associated diffraction intensity.

**Results:** Cetuximab–XRD correlations were negative in all 12 mice at every evaluated binning scale (100–400 µm; all adjusted P = 0.0039). Cohort-average correlations were more negative for cetuximab than ABY-029 at all four scales (paired adjusted P = 0.0137–0.0381). At 400 µm, mean correlations were −0.502 and −0.249, respectively. ABY-029 correlations did not differ significantly from zero after adjustment. Vehicle controls showed positive background fluorescence–XRD associations.

**Conclusions:** Cetuximab exhibited consistent inverse spatial associations with collagen-associated diffraction and stronger cohort-average associations than ABY-029. These findings show that agents targeting the same receptor can differ in their relationships with local tissue structure and support spatially resolved assessment of tissue access when selecting molecular formats for imaging and therapy.

## Introduction

The extracellular matrix (ECM) can contribute to treatment resistance in solid tumors by restricting interstitial drug transport and compressing blood vessels, thereby limiting delivery to tumor cells [1,2]. However, overall tumor accumulation does not establish that a targeted agent reaches all target-expressing regions. This distinction has therapeutic consequences: for example, improving the intratumoral penetration of an antibody–drug conjugate increased efficacy at the same administered payload dose in a mouse xenograft model [3]. Understanding how local tissue structure relates to agent distribution could therefore inform both the selection of molecular formats and interventions intended to improve tissue access [4,5]. These considerations also apply to molecular imaging, where measured uptake reflects delivery and nonspecific retention as well as target-specific binding [6]. As imaging increasingly informs targeted treatment, understanding these contributions is important for interpreting whether an imaging signal indicates adequate access by the therapeutic agent, particularly when diagnostic and therapeutic agents differ in molecular format [7,8].

Experimental studies have identified complementary roles for matrix composition, organization and agent size in tumor delivery. Measurements of macromolecular diffusion linked greater transport hindrance to collagen abundance and fibrillar organization, with differences between anatomical implantation sites demonstrating the influence of the local stromal environment [9]. Studies of fluorescent dextrans further showed that molecular weights yielding high overall tumor accumulation could nevertheless produce predominantly perivascular distributions, whereas smaller dextrans penetrated more deeply into tumor tissue [10]. Matrix-directed interventions provide additional evidence: losartan reduced collagen production and improved nanotherapeutic distribution [11], while angiotensin inhibition relieved vascular compression and enhanced chemotherapy delivery [2]. In pancreatic cancer models, enzymatic depletion of hyaluronan also restored vascular function and improved treatment response, demonstrating that delivery barriers extend beyond collagen alone [12]. More recently, spatial pharmacobiology in human pancreatic and head-and-neck tumors identified periostin-rich ECM and fibroblast neighborhoods associated with reduced therapeutic antibody delivery [13]. Complementing these structural observations, an epidermal growth factor receptor (EGFR)-targeted affibody showed greater penetration into glioma margins than a full antibody despite similar overall tumor uptake [14]. These findings motivate testing whether two co-administered agents targeting the same receptor differ in their spatial relationships with local tissue structure. Comparing an antibody and an affibody against the same tissue measurements provides a way to examine this question within a shared tumor environment.

Here, we combined two-channel fluorescence imaging with scanning synchrotron X-ray diffraction (XRD) to compare how two EGFR-targeted agents distribute relative to collagen-associated tissue structure in orthotopic AsPC-1 and PANC-1 pancreatic cancer xenografts. Cetuximab-IRDye700DX, a labeled antibody of approximately 150 kDa, and ABY-029, an IRDye800CW-labeled affibody of approximately 8 kDa [15], were co-administered, allowing their distributions to be compared within the same tumor. Diffraction intensity was reconstructed in fixed radial windows defined by a rat-tail collagen reference. This approach identifies collagen-associated features without an additional collagen-directed optical label [16]. We hypothesized that cetuximab would exhibit stronger inverse spatial associations with this signal than ABY-029. The study evaluates differences between molecular formats across spatial scales while recognizing that the agents differ in binding and pharmacokinetics as well as size.

## Materials and methods

### Study design and animals

Widefield fluorescence images and scanning XRD maps were acquired from orthotopic AsPC-1 and PANC-1 pancreatic-tail tumors in athymic nude mice. Each mouse bore one tumor, from which two slices were collected. Tumor growth was monitored by abdominal palpation over 3–4 weeks before imaging-agent administration. The analyzed cohort comprised 21 slices from 12 agent-injected mice and four slices from two vehicle-control mice. IF-10-1 was excluded because conflicting acquisition-coordinate records prevented reliable registration; IF-10-2 remained eligible.

All animal procedures, including anesthesia, postoperative analgesia and euthanasia, followed a protocol approved by the Illinois Institute of Technology Institutional Animal Care and Use Committee. NU/J athymic nude mice (The Jackson Laboratory, stock 002019) arrived at 10–12 weeks of age and acclimated for two weeks before implantation at approximately 12–14 weeks. Male and female mice were used in equal numbers. Agent-injected mice were randomly assigned to 3 h or 24 h harvest groups with a 1:1 male:female ratio. AsPC-1 and PANC-1 cells were purchased from ATCC and cultured in RPMI-1640 and high-glucose DMEM, respectively, each supplemented with 10% fetal bovine serum and 1% penicillin–streptomycin. Cryopreserved cell-line stocks underwent annual mycoplasma testing. Each mouse received 5 × 10⁵ AsPC-1 or PANC-1 cells in 50 µL of a 1:1 mixture of the corresponding culture medium and Matrigel, implanted in the pancreatic tail using a prechilled 28-gauge needle and 1 mL syringe. Cells, inoculum and all contacting supplies were kept on ice until implantation. The sample size was determined by the availability of specimens with matched fluorescence and diffraction data.

### Imaging agents and tissue preparation

Cetuximab (Eli Lilly and Company, Indianapolis, IN) was conjugated in house to IRDye700DX NHS ester (LICOR Biosciences, Lincoln, NE) and purified using centrifugal ultrafiltration with a 10 kDa molecular-weight cutoff. ABY-029, an IRDye800CW-labeled EGFR-targeted affibody, was provided by the laboratory of Kimberley S. Samkoe at Dartmouth College [17]. Each agent-injected mouse received 0.1 nmol cetuximab-IRDye700DX and 0.1 nmol ABY-029 together in 50 µL sterile 1× phosphate-buffered saline (PBS) by tail-vein injection. Vehicle-control mice received 50 µL sterile 1× PBS without either imaging agent. Separate agent-injected cohorts were harvested at 3 or 24 h.

Excised tumors were bisected, embedded in optimal cutting temperature compound and immersed in a dry ice–isopentane slurry for 60 s. The frozen blocks were removed with tweezers, held on dry ice and stored at −80 °C until sectioning. A Leica cryostat produced 500 µm slices, which were mounted directly onto chilled Ultralene sheets in photographic slide mounts. Mounted slices were stored in a chilled cassette case at −80 °C.

Mounted slices were transported on dry ice for Pearl imaging at Illinois Institute of Technology. They spent approximately 15 s in the imager before returning to dry ice and then −80 °C storage. On the day of XRD acquisition, slices were transported on dry ice to BioCAT at the Advanced Photon Source, Argonne National Laboratory, and scanned in a temperature- and pressure-controlled chamber. After scanning, samples returned to dry ice and then long-term storage at −80 °C. A cross on each mount served as the positional reference; stage coordinates were recorded with the beam aligned to the cross.

### Fluorescence and diffraction acquisition

The Pearl near-infrared imaging system (LI-COR) acquired white-light and 700 nm and 800 nm fluorescence images at 85 µm pixel spacing. Cetuximab-IRDye700DX was measured in the 700 nm channel and ABY-029 in the 800 nm channel. XRD was acquired at BioCAT on a nominal 100 µm raster. Image acquisition was done using a PILATUS3 1M detector (981 × 1,043 pixels; 172 µm detector-pixel pitch) and 1.0 s exposure per position. Rat-tail collagen reference and tumor scans used the same acquisition geometry.

### Reconstruction of collagen-associated XRD maps

Collagen-associated XRD denotes the summed diffraction intensity in three radial windows selected from a rat-tail collagen positive control reference. Original detector images were azimuthally averaged using pyFAI 2024.1.0. A local baseline was estimated with the convex-envelope procedure implemented in MuscleX 2.1.1 [16], and the baseline-subtracted profile was integrated within each window using Simpson integration. The three areas were summed at each scan position and mapped using its recorded raster coordinates. The saved detector mask excluded the beamstop and other masked pixels. No blank-image, mounting-film or fluorescence-based subtraction was applied. The same windows and processing settings were used for every tumor slice. Reference windows were chosen from rat-tail patterns without using fluorescence intensities or fluorescence–XRD correlations.

### Registration and tissue selection

Stage coordinates were mapped to Pearl pixels using the mount reference, pixel spacing and stage-axis convention. Mount edges informed rotation, and tissue boundaries and cracks guided manual translation and rotation for registration refinement. The resulting transformation was saved for each slice. Contrast enhancement assisted landmark inspection but did not alter quantitative intensities.

Freehand tissue masks were saved in the fluorescence frame. Bilinear interpolation at 29 points approximating the circular sampling footprint was averaged with equal weighting to obtain one value per channel at each XRD position. Nearest-neighbor mask sampling required every footprint point to lie within tissue. Positions were retained only when both fluorescence values and collagen-associated XRD intensity were finite. Tissue was selected anatomically without a correlation-based intensity threshold. Reviewed control masks retained all acquired positions. No additional fluorescence background subtraction was applied.

### Binning and correlation estimates

The native grid was divided into nonoverlapping 1×1, 2×2, 3×3 or 4×4 blocks, corresponding to nominal sides of 100, 200, 300 or 400 µm. A block was eligible only if all expected raster positions had been acquired and at least 75% met the tissue-validity criteria, with the required number rounded up to the next integer. Both fluorescence channels and collagen-associated XRD were averaged over the same valid positions. Incomplete edge cells were discarded. Correlations required at least six paired bins and nonzero signal variation.

Pearson correlations related each fluorescence channel to collagen-associated XRD. Because Pearson correlation is invariant to a positive constant scaling factor and an additive constant, cross-channel concentration calibration was not required for comparing spatial correlations. This invariance does not account for spatially varying attenuation, background or differences in measurement noise. Slice correlations were transformed as z(r) = arctanh(r), averaged equally within mouse and then equally across mice, and back-transformed for display. Principal 95% confidence intervals used 2,000 mouse-bootstrap resamples, preserving agent pairing. Mice, rather than pixels or slices, were the biological replicates.

### Regional analysis and coarse binning

Regional analyses illustrated two AsPC-1 sections collected at 24 h. Applying the same grid rules at 800 µm (8×8) produced 13 bins and 818 matched positions in IF-9-1. In IF-12-2, excluding bin 1 because it overlapped a dark white-light region left 19 bins and 1,171 matched positions. These examples were selected to illustrate spatial variation rather than to estimate its prevalence. A separate sensitivity analysis propagated the IF-12-2 regional exclusion to the finer scales.

For each illustrated section, each agent’s bin-mean fluorescence was divided by its unweighted mean across the retained bins. Ordinary least-squares lines and Pearson correlations describe within-specimen variation; no bin-level significance test was performed.

The same 800 µm rules were applied to all included slices, retaining the IF-12-2 exclusion. Eligible slices required at least six bins. Mouse-level paired differences used exact two-sided sign flips and a separate 20-test Holm family covering bin sizes from 100 to 2,000 µm in 100 µm increments. Scales with fewer than three eligible mice were not tested and retained a P = 1 slot in this family.

### Statistical comparisons

For each spatial scale, the principal effect was the paired difference between mouse-level Fisher-transformed correlations, Δz = z̄₇₀₀ − z̄₈₀₀; negative values indicate a more negative correlation for cetuximab than for ABY-029. Exact two-sided sign-flip tests enumerated all 4,096 assignments for the 12 mouse differences, assuming independent mice and sign symmetry under the null.

Holm correction covered the four spatial-scale comparisons [19]. In a separate family, each agent’s mean mouse-level Fisher-z correlation was tested against zero by the same two-sided procedure, with Holm correction across two agents and four scales (eight tests). Both raw and adjusted P values are reported. Confidence intervals are pointwise and are not multiplicity-adjusted tests.

Time comparisons within cell line and cell-line comparisons within time used exact two-sided label permutations, assuming exchangeability under the null. Separate 24-test Holm families covered two strata, three outcomes and four scales. The outcomes were the cetuximab–XRD correlation, ABY-029–XRD correlation and their paired Fisher-z difference. The 200 µm subgroup display uses the same mouse estimates as the principal analysis.

The vehicle-control references were HQ-1 (PANC-1) and HQ-2 (AsPC-1), each with two slices from one orthotopic pancreatic-tail tumor. They used the same reviewed tissue-mask, beam-footprint, binning and within-mouse averaging rules as agent-injected tumors. All 1,164 control positions were retained. With one control mouse per cell line, comparisons were descriptive.

### Supporting analyses and reproducibility

Supporting analyses assessed registration, tissue selection, blur and within-mouse weighting. Registration scenarios were compared on tissue valid under every scenario.

MATLAB R2024b handled spatial export and mask/registration inputs. Diffraction reconstruction used MuscleX 2.1.1 and pyFAI 2024.1.0. Statistical analysis used Python with NumPy 1.26.4, pandas 2.2.0 and SciPy 1.11.4; figures used Matplotlib 3.9.2. OpenAI Codex was used to consolidate pre-existing analysis code.

## Results

### Paired fluorescence and XRD measurements

Cetuximab had negative collagen-associated XRD correlations in all 12 mice at every principal spatial scale. Cohort-average correlations were also more negative for cetuximab than ABY-029 at all four scales, with each paired comparison remaining significant after Holm adjustment. The agent-injected cohort comprised 21 slices from 12 mice: four AsPC-1 mice at each harvest time and two PANC-1 mice at each time.

Fluorescence and XRD were measured from the same mounted slices (Figure 1). Figure 2 shows spatially matched white-light, cetuximab, ABY-029 and collagen-associated XRD images from IF-12-1 (AsPC-1, 24 h; A–D) and IF-6-3 (PANC-1, 3 h; E–H). These fields illustrate regional variation within individual sections; their different harvest times preclude a direct cell-line comparison.

**Figure 1.**
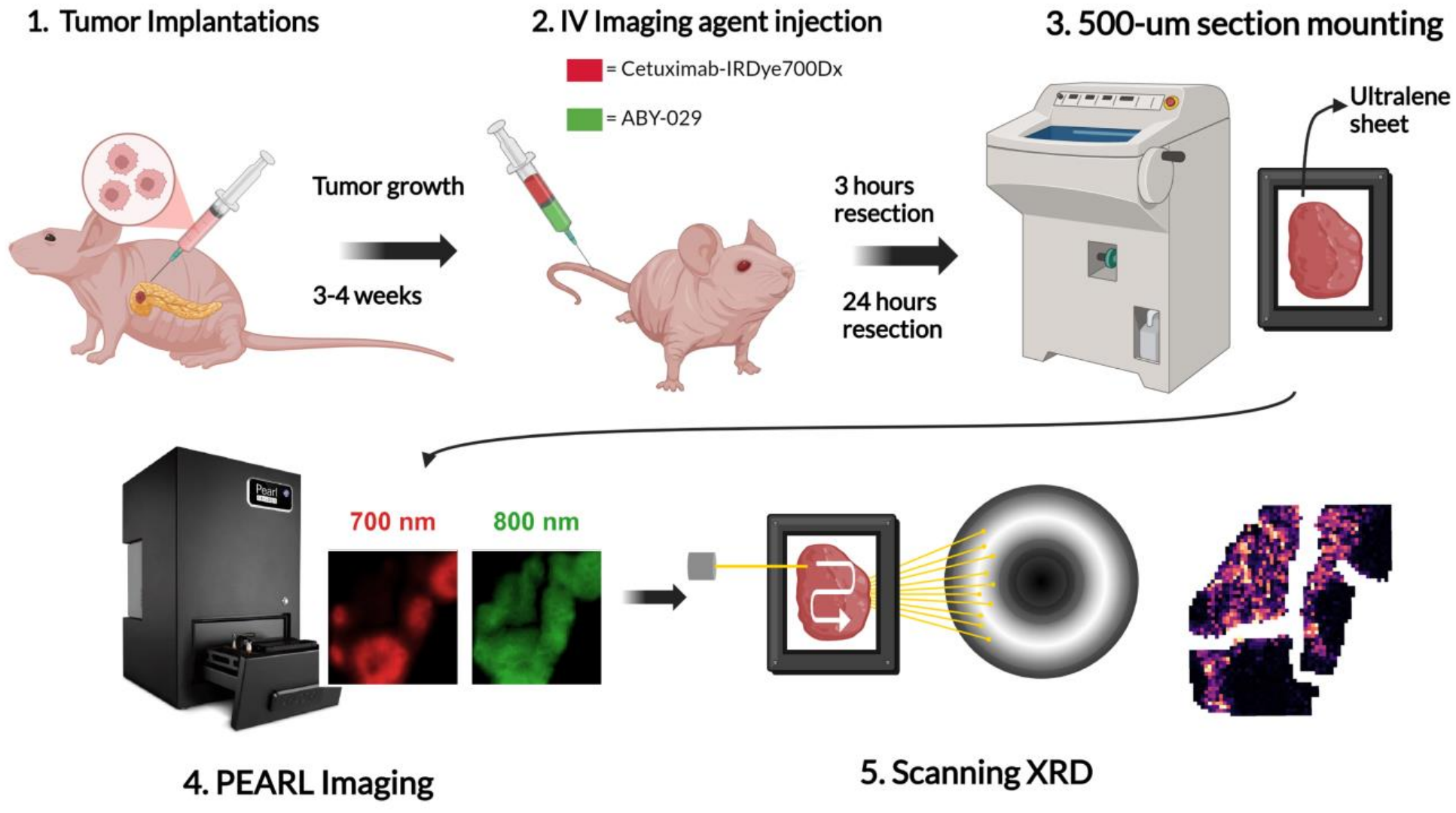
Experimental workflow. Orthotopic pancreatic-tail tumors received co-administered cetuximab-IRDye700DX and ABY-029 and were harvested at 3 or 24 h. Mounted 500 µm slices underwent Pearl fluorescence imaging and scanning synchrotron XRD. The fluorescence insets show cetuximab at 700 nm (red) and ABY-029 at 800 nm (green); the collagen-associated XRD map uses magma. Measured insets are from IF-12-2 and use independent display limits. Diffraction rings are illustrative and do not define the integration windows. Workflow artwork was provided by the authors as a BioRender illustration. XRD, X-ray diffraction.

**Figure 2.**
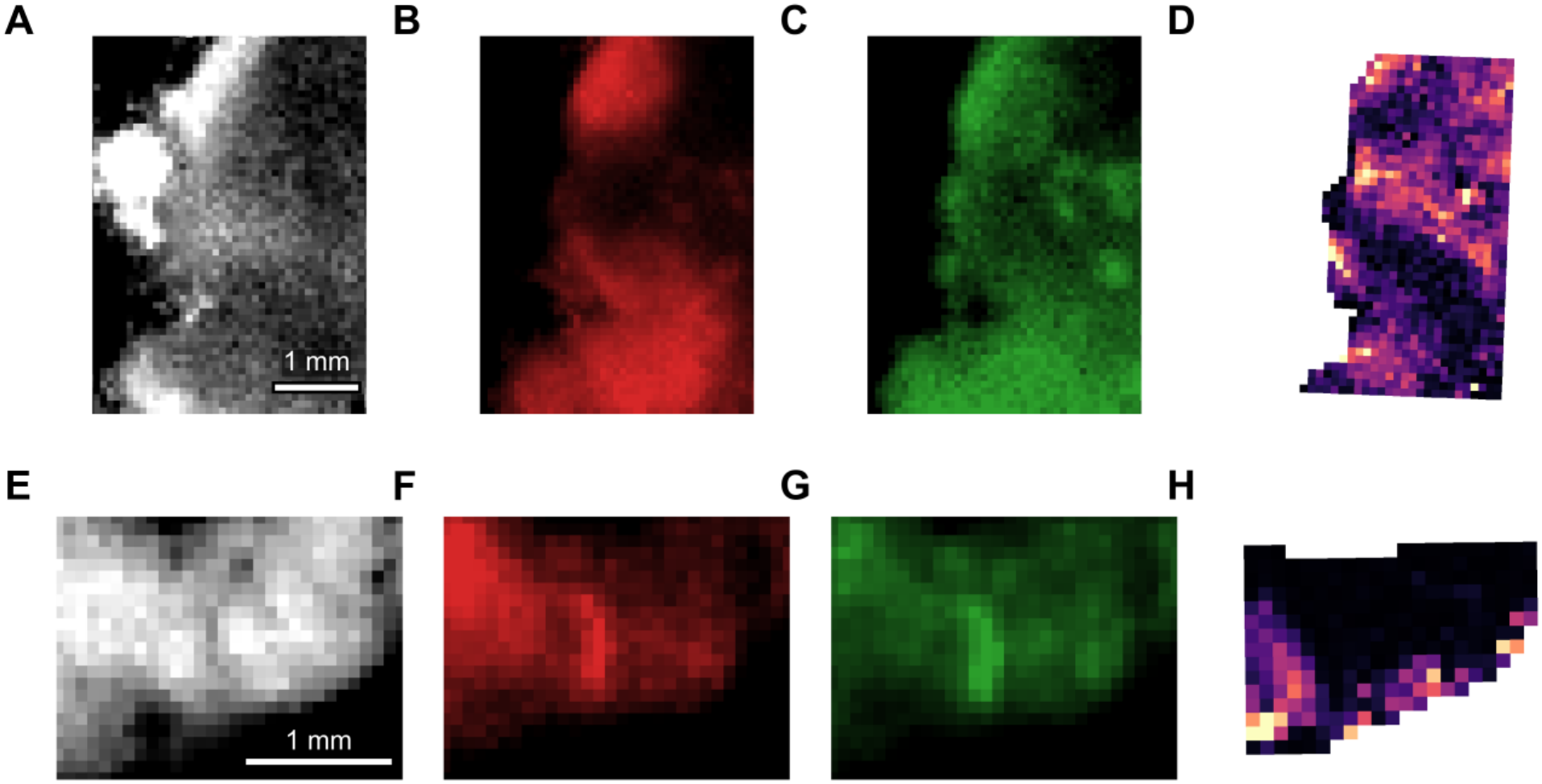
Spatially matched fluorescence and collagen-associated XRD. A–D, IF-12-1 (AsPC-1, 24 h); E–H, IF-6-3 (PANC-1, 3 h). Columns show white light, cetuximab fluorescence (red), ABY-029 fluorescence (green) and collagen-associated XRD (magma). XRD maps are restricted to the tissue selection. Scale bars, 1 mm. Independent display limits emphasize spatial patterns and do not permit cross-agent or cross-sample concentration comparisons. The examples illustrate the modalities at different harvest times rather than a controlled cell-line comparison. XRD, X-ray diffraction.

The tissue-based analysis retained 15,445 native positions. All 12 agent-injected mice contributed at every principal scale; all 21 slices contributed at 100 and 200 µm, and 20 at 300 and 400 µm.

### Regional analyses reveal differences in the agents’ spatial associations

Both AsPC-1 sections harvested at 24 h showed a strong inverse cetuximab–collagen-associated XRD association (Figure 3). In IF-9-1, the 13 retained 800 µm bins yielded r = −0.871 for cetuximab and −0.368 for ABY-029 (Figure 3E). Regions with stronger collagen-associated diffraction therefore had lower cetuximab fluorescence, while the ABY-029 distribution varied less consistently with this signal.

**Figure 3.**
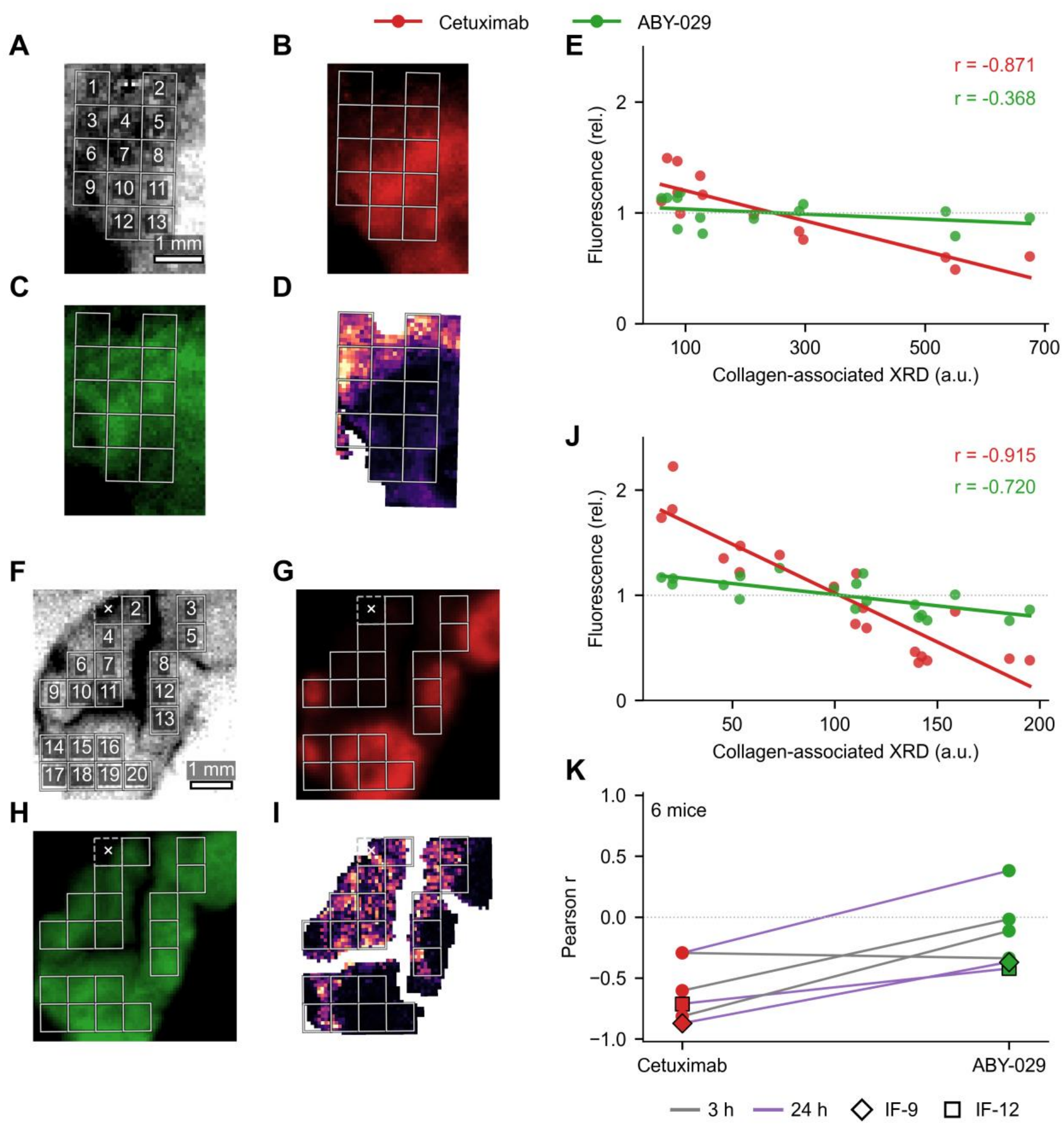
Regional collagen-associated XRD associations in two AsPC-1 tumors and the cohort eligible for 800 µm binning. A–D, IF-9-1; F–I, IF-12-2, both harvested at 24 h. Each 2×2 image block shows white light, cetuximab fluorescence, ABY-029 fluorescence and collagen-associated XRD in panel order, with 800 µm bins outlined. IF-9-1 retains 13 bins and 818 positions; IF-12-2 retains 19 bins and 1,171 positions after excluding bin 1 (×) because it overlapped a dark region in the white-light image. Scale bars, 1 mm. E, J, Each agent’s bin-mean fluorescence is divided by its mean across retained bins; lines are descriptive linear fits and annotations give Pearson r. K, Paired mouse-level correlations for all six eligible AsPC-1 mice (seven slices), with section correlations averaged within mouse on the Fisher-z scale and back-transformed. Gray and purple connecting lines indicate 3 h and 24 h, respectively; diamonds and squares identify IF-9 and IF-12. The two-sided paired sign-flip test gives raw P = 0.0625 and Holm-adjusted P = 0.8750 across 20 scales. Image display limits are independent. a.u., arbitrary units; rel., relative; XRD, X-ray diffraction.

In IF-12-2, the 19 retained bins showed inverse correlations for both agents, with a stronger relationship for cetuximab (r = −0.915) than ABY-029 (r = −0.720; Figure 3J). These examples demonstrate that a weaker inverse association for the smaller agent can coexist with different degrees of shared spatial variation. The selected fields illustrate the relationship; cohort-level inference is based on mice.

At 800 µm, seven slices from six AsPC-1 mice retained sufficient bins; no PANC-1 slice was eligible. Back-transformed mouse-level mean correlations were −0.652 for cetuximab and −0.152 for ABY-029 (Figure 3K). Paired Δz was −0.625, with lower cetuximab correlations in five of six mice. The paired difference did not reach significance at this coarse scale (raw P = 0.0625; Holm-adjusted P = 0.8750).

### Cetuximab shows consistent inverse associations and stronger paired differences

Mean cetuximab–collagen-associated XRD correlations were −0.295, −0.393, −0.421 and −0.502 at 100, 200, 300 and 400 µm, respectively (Figure 4A). All 12 mouse estimates were negative at every scale. Two-sided tests against zero were significant throughout (raw P = 0.000488; Holm-adjusted P = 0.0039 at each scale). Corresponding ABY-029 mean correlations were −0.146, −0.216, −0.218 and −0.249; none differed from zero after adjustment (adjusted P = 0.2798, 0.2798, 0.2798 and 0.1699). This does not establish an absence of an ABY-029 association.

**Figure 4.**
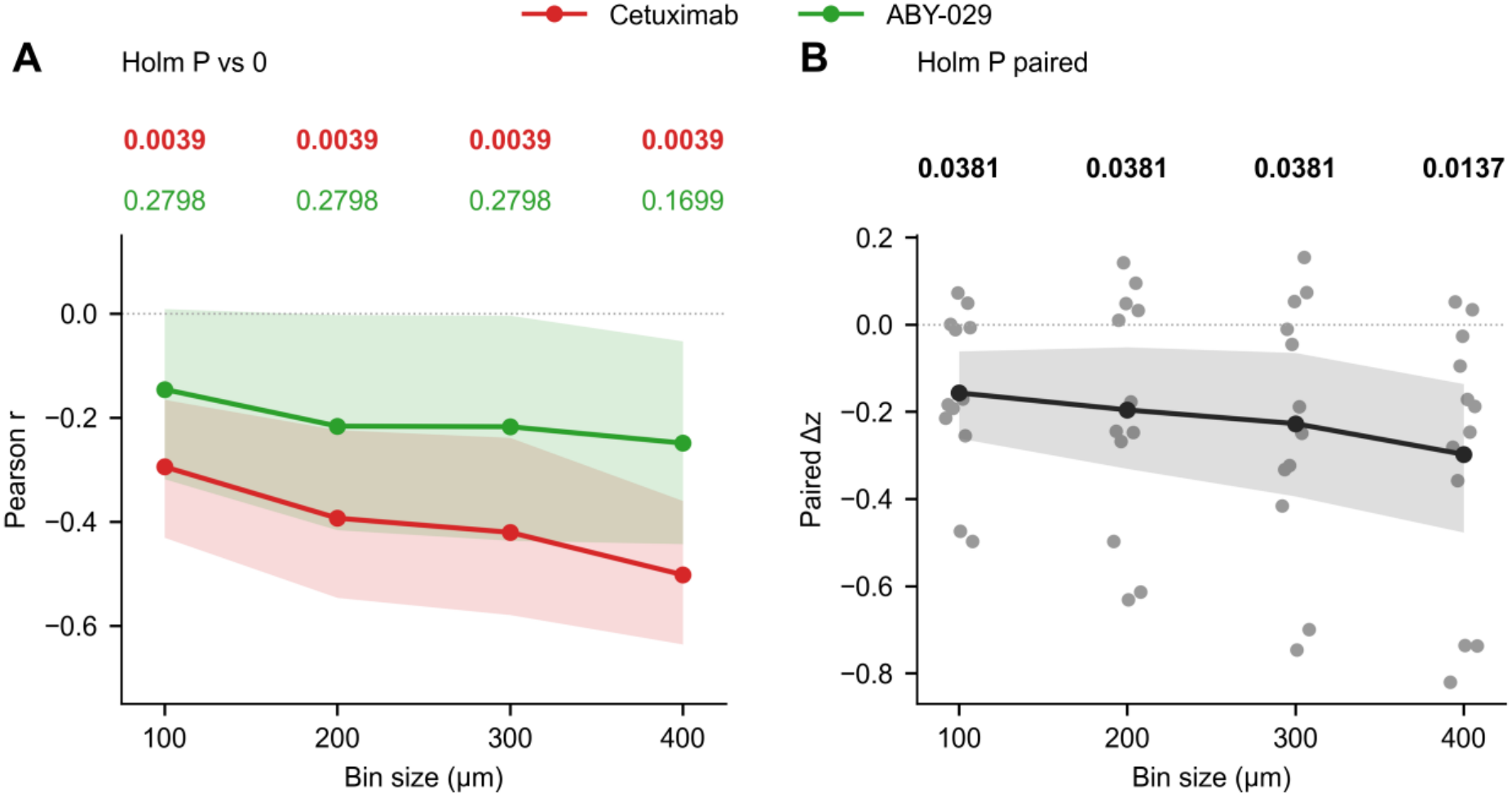
Cetuximab has consistent negative collagen-associated XRD correlations and more negative correlations than ABY-029. A, Section correlations are Fisher-z averaged within mouse, then equally across 12 mice and back-transformed. Colored P values test each agent’s mean mouse-level Fisher-z correlation against zero using two-sided exact sign flips and Holm correction across eight agent-by-scale tests. B, Paired Δz = z̄₇₀₀ − z̄₈₀₀; negative values indicate a more inverse cetuximab association. Gray dots are individual mice. Black P values use paired two-sided exact sign flips and Holm correction across four scales. Bold values are below 0.05. Shading shows pointwise 95% intervals from 2,000 mouse-bootstrap resamples. All 21 slices contribute at 100 and 200 µm; 20 contribute at 300 and 400 µm. The tests assume independent mice and null sign symmetry. Interval overlap between agents is not a test of their paired difference. XRD, X-ray diffraction.

The paired agent difference was negative at every scale (Δz = −0.157, −0.196, −0.227 and −0.298), with Holm-adjusted P values of 0.0381, 0.0381, 0.0381 and 0.0137, respectively (Figure 4B). At 400 µm, the 95% CI for Δz was −0.477 to −0.136. Cetuximab correlations were lower than ABY-029 correlations in 9, 7, 9 and 10 of 12 mice across the four scales. The paired tests directly establish a difference between the agents, independently of whether either agent’s correlation differs from zero.

### Paired agent differences across cell lines and harvest times

At 200 µm, mean paired differences were negative in AsPC-1 tumors at 3 h and 24 h (Δz = −0.278 and −0.389), but positive in the two PANC-1 mice at each time (+0.064 and +0.095; Figure 5). Thus, the cohort-average paired difference was not uniform across tumor models. Exact subgroup comparisons did not detect time or cell-line differences after adjustment (all Holm-adjusted P = 1.00). The small subgroup sizes, particularly for PANC-1, limit interpretation of these null tests.

**Figure 5.**
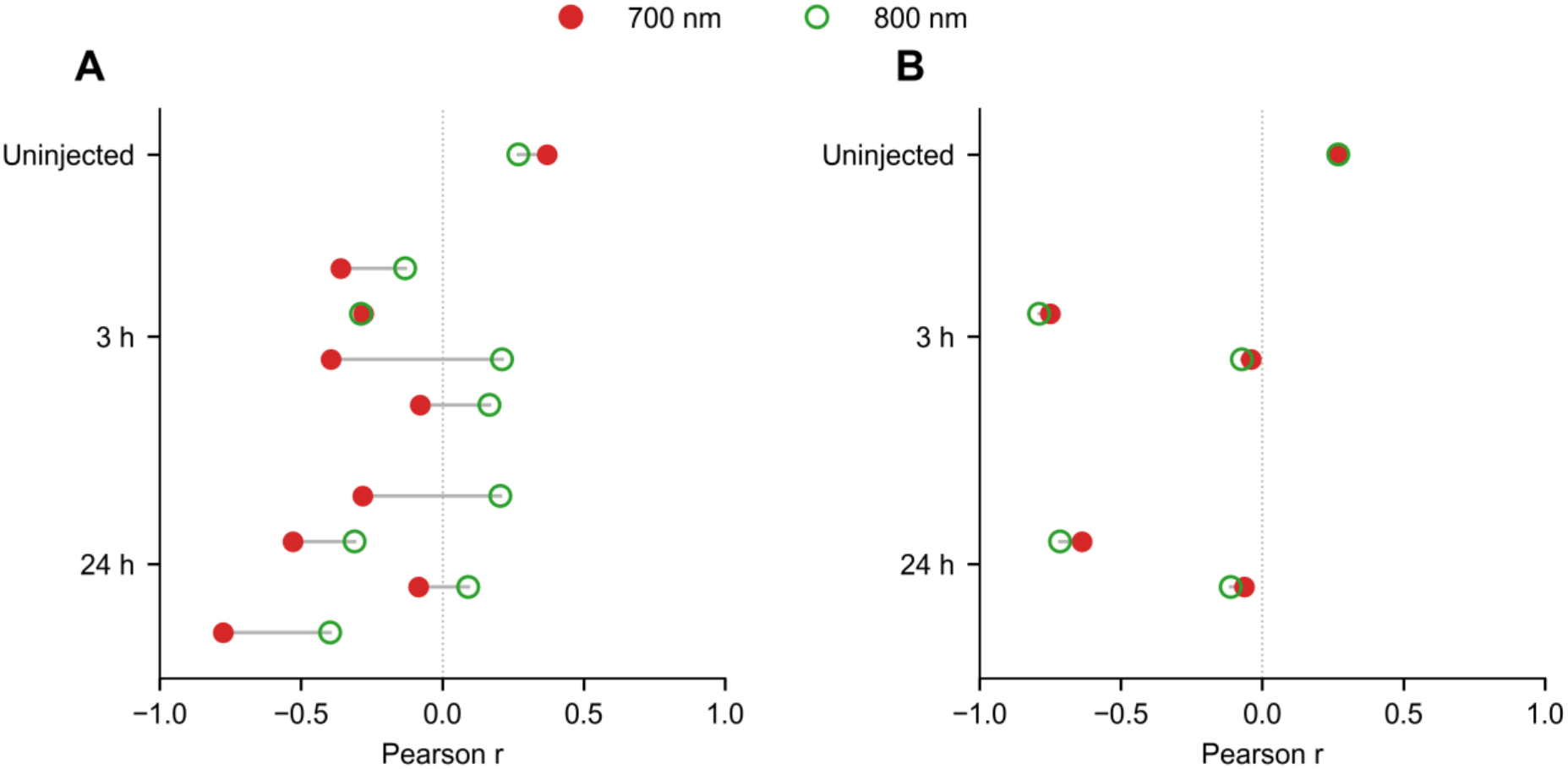
Mouse-level collagen-associated XRD correlations by tumor model and harvest time. A, AsPC-1; B, PANC-1. Each connected pair is one mouse at 200 µm binning, with section correlations averaged on the Fisher-z scale. Filled red points show the 700 nm channel; open green points show the 800 nm channel. There are four injected AsPC-1 mice and two injected PANC-1 mice per timepoint. Vehicle controls comprise one mouse per cell line, each with two reviewed slices. Control fluorescence represents background rather than tracer delivery. No background subtraction or autofluorescence calibration is applied. XRD, X-ray diffraction.

### Vehicle-control tumors show positive background fluorescence–XRD associations

Both vehicle-control tumors showed positive fluorescence–collagen-associated XRD correlations in both channels (Figure 5). At 200 µm, 700 nm and 800 nm correlations were +0.270 and +0.269 in PANC-1 control HQ-1 and +0.370 and +0.269 in AsPC-1 control HQ-2. These associations were present without administered imaging agents, contrasting with the negative cohort-average correlations in agent-injected tumors.

## Discussion

Cetuximab showed a consistent inverse spatial relationship with collagen-associated XRD across orthotopic pancreatic tumor specimens: every mouse had a negative association at each of the four principal scales. The co-administered ABY-029 had less negative cohort-average correlations, and paired comparisons supported this difference at every scale after correction.

The result concerns where the agents accumulated within the same tumors, rather than their whole-tumor uptake or calibrated concentration. It is consistent with more restricted local cetuximab accumulation in regions with stronger collagen-associated diffraction. The two-sided tests against zero establish the consistency of the inverse cetuximab relationship; the paired tests separately establish the difference between agents. An insignificant ABY-029 test against zero does not establish that the affibody is unaffected by tissue structure.

The stronger inverse association of cetuximab with collagen-associated XRD is consistent with more restricted local accumulation. Previous measurements linked impaired macromolecular diffusion to collagen abundance and organization, with stronger hindrance for larger probes [9]. Second-harmonic generation (SHG) imaging subsequently connected fibrillar collagen structure with diffusive transport and its modification by relaxin [20]. Studies combining SHG with fluorescence imaging also demonstrated different effects of matrix-degrading enzymes on immunoglobulin G (IgG) and liposomal doxorubicin delivery [21]. In pancreatic tumors, regional collagen measurements have been linked to tissue pressure, vascular patency and verteporfin delivery [22]. Our study extends these observations by comparing two co-administered agents targeting the same receptor against shared local structural measurements, revealing a more negative association for the antibody than for the affibody. These findings also complement the observation that an EGFR-targeted affibody penetrated glioma margins more effectively than cetuximab despite similar overall tumor uptake [14]. Molecular size, however, operates alongside binding and systemic exposure: models of tumor targeting describe trade-offs among extravasation, tissue transport, affinity and clearance [5]. Binding-site barriers can restrict penetration through perivascular binding [23], while Krogh-cylinder simulations predict more uniform access to distant binding sites by smaller targeting constructs [24]. Although binding-site barriers were not evaluated here, the observed relationships with collagen-associated diffraction motivate studying these processes together. Combining spatial measurements of matrix structure, vascular proximity, and target expression with transport–binding models could help determine how local matrix conditions and binding jointly influence tissue access.

The diffraction measurement links the paired fluorescence distributions to collagen-associated diffraction features. The same radial windows were used for all specimens, providing a consistent structural measurement. Their intensities, however, depend on orientation, ordering and sampled material as well as collagen abundance [16,18]. Without independent spacing calibration and complementary tissue characterization, the signal cannot be attributed exclusively to collagen or used to distinguish collagen content from organization. This distinction matters biologically: human spatial pharmacobiology studies associate reduced antibody delivery with particular ECM and fibroblast neighborhoods [13], while earlier osteosarcoma studies found no simple relationship between IgG uptake and bulk collagen content [25]. Mechanical imaging provides a complementary perspective; elastography studies in pancreatic xenografts link greater local stiffness to reduced drug delivery and vascular patency, with stiffness also associated with collagen density [26,27]. Combining diffraction with histology and mechanical measurements could help distinguish how matrix composition, organization and mechanics relate to agent distribution.

The regional examples and multiscale results support the same overall observation while showing its heterogeneity. Both illustrated sections had stronger inverse cetuximab correlations, but ABY-029 was moderately associated with collagen-associated XRD in IF-9-1 and strongly associated in IF-12-2. The cohort-average paired difference remained negative from 100 to 400 µm. Extending the analysis to 800 µm preserved that direction, but only six AsPC-1 mice retained sufficient bins and the paired test was not significant. Coarser bins summarize broader regional variation while reducing within-slice observations and cohort eligibility; they do not identify a characteristic biological transport length. Spatially resolved measurements can therefore reveal differences obscured by a single whole-tumor uptake value [28], but their scale and support must remain explicit.

The paired fluorescence measurements allow comparison of spatial associations without requiring the two channels to share an absolute concentration scale. Pearson correlations are unaffected by a positive constant scaling factor or an additive constant, making them suitable for comparing how each agent’s signal varies with local XRD intensity despite differences in fluorophore brightness and channel sensitivity. Spatially varying background, however, can influence these relationships. Both vehicle-control tumors showed positive fluorescence–XRD correlations in both channels, demonstrating that tissue-associated background can co-vary with diffraction intensity even without administered agents. This pattern contrasts with the negative cohort-average correlations in agent-injected tumors. If a comparable positively associated background contributed to the injected specimens, it would oppose the negative agent–XRD covariance rather than generate it. Nevertheless, the magnitude and spatial variability of background may differ between channels, and one control mouse per cell line provides only a descriptive reference. The principal finding is therefore a difference in spatial association, rather than a comparison of absolute agent concentrations. Future measurements combining matched background characterization and fluorescence calibration could extend this approach to quantify local agent abundance and assess the contribution of wavelength-dependent optical sampling.

Co-administration enabled within-tumor comparisons, with mice treated as independent biological replicates. The cohort-average paired difference remained negative across all 147 tested registration scenarios at each principal scale. Paired comparisons on tissue common to every scenario did not pass multiplicity adjustment, indicating that statistical strength depends on spatial support. Registration uncertainty was not independently quantified. The paired differences also varied by tumor model: at 200 µm, AsPC-1 group means favored a more negative cetuximab association, whereas PANC-1 group means had the opposite sign. Small subgroup sizes limit the ability to distinguish tumor-model and time effects. The 3 h and 24 h measurements capture agents with different transport, binding and clearance kinetics [5]; equal injected molar doses do not place them at equivalent stages of tissue uptake. Additional cohorts, independent registration assessment and more harvest times would help establish the generality and temporal evolution of the relationship.

These findings have implications for both the selection of targeted agents and strategies intended to improve their access to tumor tissue. When diagnostic and therapeutic agents differ in molecular format, their shared target does not necessarily imply equivalent access to target-expressing regions. The clinical evaluation of ABY-029 for fluorescence imaging in soft-tissue sarcoma highlights the translational relevance of understanding how smaller targeting agents distribute within heterogeneous tumors [15]. Preclinical studies of losartan have demonstrated improved nanotherapeutic distribution following reduced collagen production and improved chemotherapy delivery through relief of vascular compression [2,11]. Complementary studies targeting collagen crosslinking with pan-lysyl oxidase inhibition have reduced stromal remodeling, improved tumor perfusion, and enhanced gemcitabine response in pancreatic cancer models [29]. Applying paired fluorescence–XRD measurements to such interventions could reveal whether changes in collagen-associated diffraction features accompany improved agent accumulation in previously poorly accessed regions. Coupling these measurements with target expression and treatment response would help connect structural changes to their therapeutic consequences. MR elastography, which can detect collagen-associated differences in tumor mechanics and their response to stromal modification, offers a complementary approach for investigating the mechanical consequences of matrix remodeling noninvasively [30]. The broader opportunity is to evaluate targeted delivery not only by how much agent accumulates in a tumor, but also by where it accumulates and which tissue environments remain difficult to reach.

## Conclusions

Cetuximab showed negative collagen-associated XRD correlations in every analyzed mouse and stronger inverse cohort-average associations than ABY-029 across 100–400 µm scales. Spatially matched fluorescence and diffraction thus distinguished the distributions of two co-administered agents within the same tumors. These findings support further investigation of collagen-associated tissue structure as a determinant of local agent access, while separating spatial association from collagen concentration and causal transport effects.

## Abbreviations

AI: artificial intelligence
ATCC: American Type Culture Collection
CI: confidence interval
DMEM: Dulbecco’s modified Eagle medium
ECM: extracellular matrix
EGFR: epidermal growth factor receptor
IgG: immunoglobulin G
MR: magnetic resonance
NHS: N-hydroxysuccinimide
PBS: phosphate-buffered saline
SHG: second-harmonic generation
XRD: X-ray diffraction.

## Acknowledgements

This research used BioCAT resources at the Advanced Photon Source, a US Department of Energy Office of Science User Facility operated by Argonne National Laboratory under contract DE-AC02-06CH11357. BioCAT was supported by National Institute of General Medical Sciences grant P41GM103622. The PILATUS3 1M detector was supported by National Institutes of Health grant S10OD018090.

OpenAI Codex assisted with consolidation of previously written analysis code. The authors are responsible for the accuracy and integrity of the AI-assisted content.

## Ethics approval

Animal procedures were approved by the Illinois Institute of Technology Institutional Animal Care and Use Committee.

## Data and code availability

The data and code used in this analysis are available on request from the corresponding author, Kenneth M. Tichauer.

